# Elevation drives activity of soil bacteria, but not of bacterial viruses

**DOI:** 10.1101/2021.12.14.472558

**Authors:** D. Merges, Alexandra Schmidt, Imke Schmitt, Eike Lena Neuschulz, Francesco Dal Grande, Miklós Bálint

## Abstract

Soil microbial diversity affects ecosystem functioning and global biogeochemical cycles. Soil bacterial communities catalyze a diversity of biogeochemical reactions and have thus sparked considerable scientific interest. One driver of bacterial community dynamics in natural ecosystems has so far been largely neglected: the predator-prey interactions between bacterial viruses (bacteriophages) and bacteria. To generate ground level knowledge on environmental drivers of these particular predator-prey dynamics we propose an activity-based ecological framework to simultaneous capture community dynamics of bacteria and bacteriophages in soils. An ecological framework and specifically the analyses of community dynamics across latitudinal and altitudinal gradients have been widely used in ecology to understand community-wide responses of innumerable taxa to environmental change, in particular to climate. Here, we tested the hypothesis that the activity of bacteria and bacteriophages co-decline across an elevational gradient. We used metatranscriptomics to investigate bacterial and bacteriophage activity patterns at 5 sites across 400 elevational meters in the Swiss Alps in 2015 and 2017. We found that metabolic activity (transcription levels) of bacteria declined significantly with increasing elevation, but activity of bacteriophages did not. We showed that bacteriophages are consistently active in soil along the entire gradient. Bacteriophage activity pattern, however, is divergent from that of their putative bacterial prey. Future efforts will be necessary to link the environment-activity relationship to predator-prey dynamics, to understand the magnitude of viral contributions to mobilize bacterial cell carbon when infection causes bacterial cell death, a process that may represent an overlooked component of soil biogeochemical cycles.

## Introduction

Soil microbiomes are key for ecosystem functioning and play pivotal roles in global biogeochemical cycles (i.e., C and N cycling) (Braga et al., 2020; Pratama & van Elsas, 2018). The soil microbiome harbors groups of highly diverse organisms, such as bacteria, archaea, fungi and protozoa, including viruses that infect them (Kimura et al., 2008; Pratama & van Elsas, 2018). Bacteria and archaea catalyze a diversity of biogeochemical reactions, many of which are climate relevant (Hallin & Bodelier, 2020; Monteux et al., 2020). Therefore, bacterial and archaeal functional assessment has sparked considerable interest (Hallin & Bodelier, 2020; Kimura et al., 2008; Pratama & van Elsas, 2018). Soil bacteria are increasingly considered in soil studies, but their respective viruses are readily neglected, resulting in a lack of understanding of how the interaction of bacteria with their viruses influences soil functioning (Ashelford et al., 2003; Kimura et al., 2008; Marsh & Wellington, 1994; Pratama & van Elsas, 2018). Viruses may impact soil communities by a) controlling microorganismal population dynamics as predators (Breitbart et al., 2018; Morella et al., 2018), and b) providing genes and functions, which may alter ecosystem properties, e.g. carbon degradation (Emerson et al., 2018; Pratama & van Elsas, 2018; Trubl et al., 2018).

Bacteriophages, or short ‘phages’, are viruses that prey on bacteria (Hoffmann et al., 2007). From the point of view of trophic interactions, the relationship between bacteria and their viruses can be regarded as that of a prey and its predator (Chao et al., 1977; Weinbauer, 2004; Weitz & Dushoff, 2008). The interaction is initiated during a random encounter of the bacterium and a phage with its adsorption to specific receptor sites on the bacterial cell (Chao et al., 1977). Subsequently, the genome of the phage is injected into the bacterium (Chao et al., 1977). From this stage on, phages can be classified as either virulent or temperate (Weitz & Dushoff, 2008). Virulent phages reproduce within the bacterial cell, kill their hosts and release an array of infective phage particles without undergoing an extended intracellular phase, whereas temperate phages can incorporate their genome into that of the host and remain dormant (Chao et al., 1977; Weinbauer, 2004; Weitz & Dushoff, 2008). Viruses may affect carbon cycling and carbon degradation in soil, 1) by mobilizing bacterial cell carbon when viral infection causes bacterial cell death and 2) by degrading plant-derived polymers into monosaccharides and oligosaccharides – which, in turn, can be metabolized by microbes resulting in CH4 and CO2 emissions – via glycoside hydrolases (Emerson et al. 2018). The full extent of bacteria-phage interactions across different environments, including soil, is poorly understood (Braga et al., 2020; Emerson et al., 2018; Roux et al., 2021; Starr et al., 2019).

The main factor limiting phage occurrence is the presence of bacterial hosts (Olszak et al., 2017; Weitz & Dushoff, 2008). Thus, all ecosystems with metabolically active bacteria are expected to have abundant and diverse phage populations (Marsh & Wellington, 1994). Accordingly, previous studies using metatranscriptomics found active phage communities when bacteria where active in soils (Emerson et al., 2018; Starr et al., 2019).

The integration of an ecological framework, such as activity analyses across gradients, into bacteria-virus interaction studies can address questions which have been mostly neglected in virology, but are key to advance our understanding on the ecosystem consequences of predator-prey dynamics (Sommers et al., 2021). Numerous previously published articles reported changes in bacterial metabolic activity across latitudinal and elevational gradients (Chase et al., 2021; Margesin et al., 2009; Ren et al., 2021; Rivkina et al., 2000; Schinner, 1982). Across these gradients, the metabolic activity of bacteria has been linked to climatic factors, such as mean annual temperature, precipitation as well as soil properties (bulk density, ammonium nitrogen, and total phosphorus) (Margesin et al., 2009; Ren et al., 2021; Rivkina et al., 2000; Schinner, 1982). Given these changes in bacterial metabolic activity, one would expect changes in phage activity, with a potential impact on bacterial populations (Marsh & Wellington, 1994). Thereby a replicated elevational gradient setup is highly suitable to generate ground level knowledge on these particular predator-prey interactions and to assess their community-wide responses to environmental change, in particular to climate. However, no study to date has investigated the activity of bacteria and their phage predators across an elevational gradient.

The aim of this study was to establish a baseline approach to simultaneously assess bacteria and bacteriophage activities and their drivers in soil. To understand community-wide responses to environmental change, in particular to climate, we utilize gradient analyses from an ecological framework (Sommers et al., 2021) to assess the population-wide activities of bacteria and phage communities across an elevational gradient in the European Alps. Elevational gradients allow the study of broad environmental conditions on a condensed geographic scale (Bergner et al., 2020; Neuschulz et al., 2018). Specifically, we employed a metatranscriptomic approach to assess how activities of soil bacteria and phages, with regard to the expressed metabolic pathways, respond to elevation. Assuming that phages require metabolically active hosts to support multiplication (Marsh & Wellington, 1994), we expected similar levels of activity in bacteria and phages in a given environment. Since bacterial metabolic activity is lower at higher altitudes (Margesin et al., 2009; Ren et al., 2021; Schinner, 1982), we expected a co-decline of bacterial and phage activity with increasing elevation.

## Material/Methods

### Study site & sampling

The study sites were located in the Central Alps in the eastern part of Switzerland in the Sertig valley (46°44’0.76”N, 9°51’3.5”E) near Davos (Merges et al., 2020; Neuschulz et al., 2018). For soil sampling of bacterial and phage communities, we sampled five elevational levels at 1850, 1900, 2000, 2100 and 2250 m a.s.l (Table S1). We conducted two sampling rounds, one in May 2015 and one in May 2017, resulting in a total of 10 soil samples (Table S1). Soil samples were taken with a 1 cm soil core sampler (Ehlert & Partner). For each soil sample, we took five 5-cm deep soil cores from a 15 × 15 cm^2^ area that we pooled and homogenized in a Ziploc bag (Merges et al., 2018). 10 g of homogenized soil were immediately transferred into a 50 mL Falcon filled with RNA preservative (LifeGuard Soil Preservation Solution, QIAGEN). The preserved soil was frozen at −80°C when brought to the lab.

### Lab & bioinformatics

RNA was extracted with RNeasy PowerSoil Total RNA Kit (QIAGEN) and deeply sequenced to 8GB depth per sample at NOVOGEN. We received consistently high yields, with a total of 30 – 40 Mio. reads per sample (NCBI Bioproject ID XXXXX). Sequences were quality filtered and trimmed of adapters using TRIMMOMATIC (Bolger et al., 2014). We assessed the quality of reads with fastQC (https://www.bioinformatics.babraham.ac.uk/projects/fastqc/).

### Assembly of bacterial contigs

Trimmed reads were assembled with TRINITY (Grabherr et al., 2013). Assembled contigs were taxonomically binned using the *last common ancestor* (LCA) algorithm of DIAMOND with the NCBI nr protein database and bacterial contigs were selected (Buchfink et al., 2014). Activity was assessed by mapping back raw reads to individual taxa bins using SALMON/deseq2 (please see below). PROKKA was used for functional annotation (Seemann, 2014).

### Assembly of viral contigs

Trimmed reads were assembled with rnaviralSPAdes (Lapidus & Korobeynikov, 2021; Nurk et al., 2017). Assembled contigs were screened for viral origin using VirSorter2 (Guo et al., 2021). Putative viral contigs were further quality controlled using CheckV (Nayfach et al., 2020). Passing contigs were taxonomically binned using the *last common ancestor* (LCA) algorithm of DIAMOND with the NCBI nr protein database (Buchfink et al., 2014). The produced dataset of viral contigs was subsetted to taxa exclusively associated with bacteria (i.e., bacteriophages). We used PROKKA for functional annotation (Seemann, 2014).

### Activity/transcript expression analyses

For transcript abundance quantification (i.e. estimated counts per transcript) we used SALMON v0.14.1 with the --gcBias flag (Patro et al., 2017). The --gcBias flag integrates the estimation of a correction factor for systematic biases frequently present in RNA-seq data (Patro et al., 2017). The tximport package (Soneson et al., 2016) was used to import the quantified data from SALMON into R v3.6.1 (R Core Team, 2019). For differential expression analysis, a DESeqDataSet was constructed from the tximport object with the DESeqDataSetFromTximport function from the DESeq2 package (Love et al., 2014). We tested differences between the two sampling years using the Likelihood ratio test (LRT) with Benjamini–Hochberg false discovery rate control as implemented in DESeq2 (Love et al., 2014). Transcripts were normalized using the “Relative Log Expression” normalization (RLE) (Love et al., 2014). We found no significantly different expressed transcripts between the two years (Benjamini-Hochberg adjusted p-values > 0.05). The plotPCA function was used to visualize similarities between temporal replicates (Figure S1). Due to absence of temporal effects on transcript expression, we tested the effect of elevation on the normalized read counts using generalized linear models (GLM) with negative binomial distribution (Venables & Ripley, 2002). To account for genome size differences, when comparing bacterial and phage activity, all available genome size information on taxa were retrieved from NCBI refseq database (n=5507) using the R/Bioconductor package biomaRt (Durinck et al., 2009). Analysis was repeated with a subset, where genome size information was available, by additionally normalizing read counts for genome size (i.e. dividing normalized read counts by the mean genome size of the respective taxa; Table S10 & S11).

## Results

### Taxonomic Diversity

#### Diversity of bacterial taxa

The retrieved 170.265 bacterial contigs spanned a diversity of 37 phyla encompassing 428 families (Figure 1). The most expressed transcripts belonged to the phyla *Proteobacteria, Actinobacteria, Firmicutes, Acidobacteria*, and *Bacteroidetes* (Table S2 & Table S5-S9).

**Figure 1:**
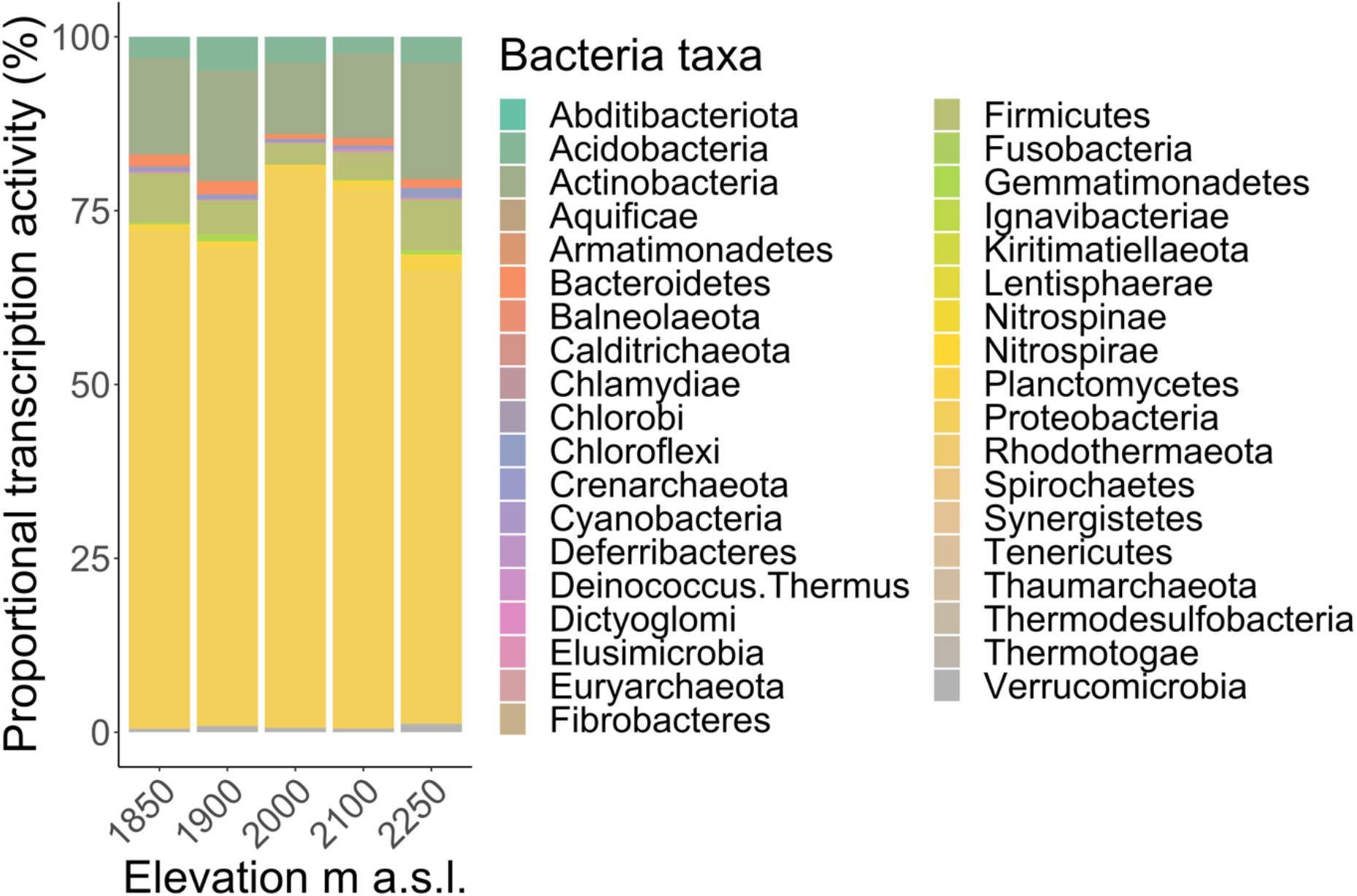
Proportional transcription activity of soil bacteria across the elevational gradient.

#### Diversity of phage taxa

We received 216 bacteriophage contigs belonging to 3 orders across 9 distinct families (Fig. 2). Of these, the double stranded DNA (dsDNA) bacterial virus families of *Autographiviridae, Demerecviridae, Herelleviridae, Myoviridae, Podoviridae, Siphoviridae* of the order *Caudovirales* showed the highest transcriptional activity (Table S3 & S4). One further viral family with DNA genomes (single stranded DNA) was detected, belonging to the family of *Microviridae* (Order *Petitvirales*), as well as one family of single stranded RNA viruses (ssRNA viruses: *Fiersviridae*, formally *Leviviridae*, Order *Norzivirales*, Table S3 & S4).

**Figure 2:**
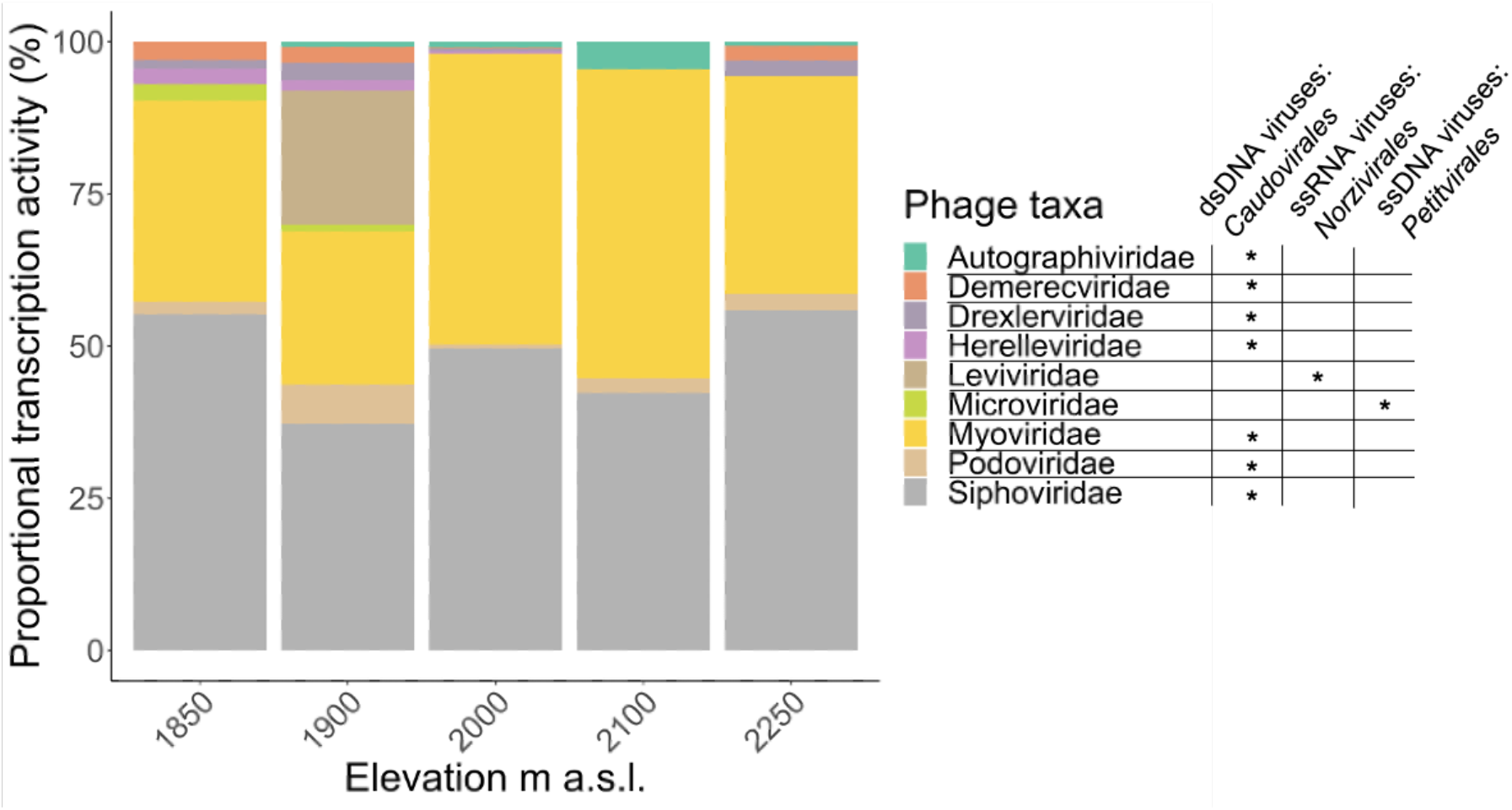
Proportional transcription activity of soil bacteriophages across the elevational gradient.

#### Bacterial and viral activity across years and elevation

Overall activity was not significantly different between the years 2015 and 2017 (p > 0.05; Fig. S1). Bacterial activity significantly declined with increasing elevation (p < 0.01, Fig. 3), whereas bacteriophage activity was not significantly affected by elevation (p > 0.05, Fig. 3). The patterns were robust when normalizing the transcriptional activity by mean genome size (Fig. S4).

**Figure 3:**
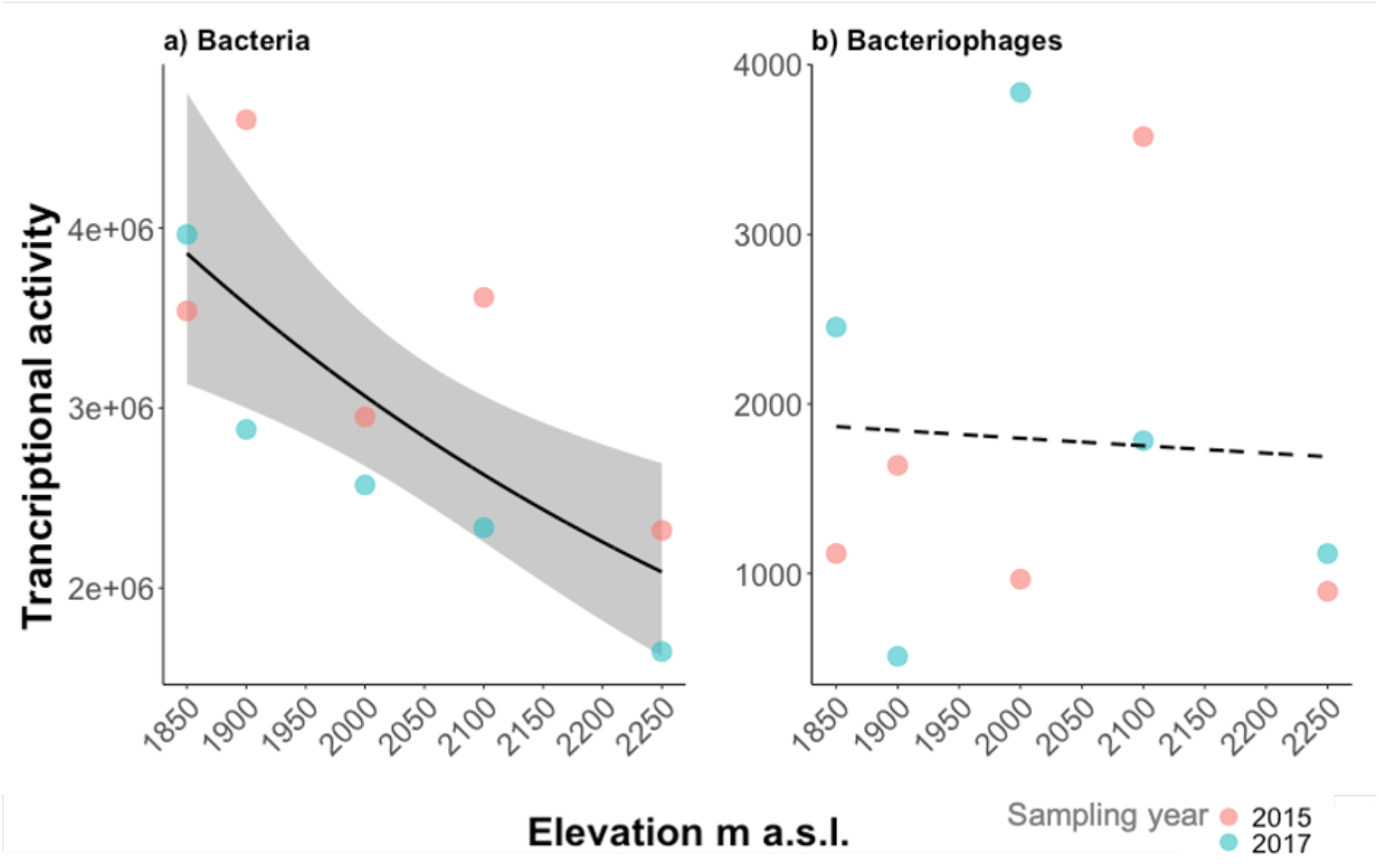
Soil bacterial (a) and phage (b) transcriptional activity across the elevational gradient. Bacterial activity (a) significantly declined with increasing elevation, whereas phage activity (b) showed no response to elevation.

#### Metabolic pathways

Annotation of expressed genes revealed a broad diversity of metabolic pathways across the bacterial taxa (Fig. S2 & S3). The majority of annotated genes were directly related to metabolic pathways, tRNA biosynthesis and the biosynthesis of secondary metabolites (Fig. S2 & S3). For viral taxa, no functional information could be retrieved. The proportional activity of metabolic pathways remained constant across the elevational gradient (Fig. S3).

## Discussion

So far, the effect of phages on soil bacterial communities has been mostly neglected (Braga et al., 2020) and only recently has been receiving interest enabled by the advance of high throughput approaches (Braga et al., 2020; Emerson et al., 2018; Trubl et al., 2018). In the present study, we applied a metatranscriptomics approach, where we identified highly diverse bacterial and phage communities in soils across an elevational gradient. In accordance with our hypothesis, bacterial metabolic activities strongly declined with increasing elevation. In contrast, we found that bacteriophages are consistently active in soil along the entire gradient.

We found the taxa *Proteobacteria, Actinobacteria, Firmicutes, Acidobacteria*, and *Bacteroidetes* to be the most metabolically active members of the soil bacterial communities (Fig. 1). Their activity strongly declined with increasing elevation (Fig. 3a). Similarly, Margesin et al. (2009) could show a decline in the relative amount these taxa as well as a decrease in activity in a gradient in Austrian Central Alps, based on measurements of soil dehydrogenase activity, with increasing elevation in alpine soils. Ren et al. (2021) reported comparable bacterial community composition and a decreasing abundance with increasing elevation, based on 16S rRNA amplicon sequencing of soils from the Qinling Mountains in central China. Both studies could link the decline in activity and abundance to lower temperatures at high elevational sites (Margesin et al., 2009; Ren et al., 2021). We suspect that temperature also underlies the pattern observed by us for bacterial transcriptional activity. This may be confirmed with a follow-up study with increased sample sizes, combined with fine-scale temperature measurements.

We detected ssRNA viruses (Family: *Leviviridae*) only in one single elevation (Fig. 2, Elevation 1900 m a.s.l., brown bar). So far, RNA phages of the *Leviviridae* family were identified in a metatransciptomics study in soils from California (Starr et al., 2019). The authors reported a generally large diversity of *Leviviridae*. However, in accordance with our findings, a heterogeneous distribution across samples and replicates (Starr et al., 2019).

We found dsDNA bacteriophages of the order *Caudovirales* to be the most dominant members of the active soil viral communities. In accordance with our study, dsDNA bacteriophages of the order *Caudovirales* are consistently reported as the most dominant viruses present in soil (Adriaenssens et al., 2017; Emerson et al., 2018; Williamson et al., 2005). (Williamson et al., 2005) showed that the majority of soil viruses in Delaware soil were bacteriophages belonging to the *Caudovirales* by combing direct counting of virus-like particles (VLPs) with morphological data gathered using TEM. Additional supporting evidence came from recent metagenomic approaches, which found members of the *Caudovirales* (specifically the families: *Myoviridae, Podoviridae* and *Siphoviridae*) made up more than 80% of the relative abundance at all sites in Antarctic soil (Adriaenssens et al., 2017). These are the families which also showed high activity in our samples (Fig. 2). A similar dominance of members of the *Caudovirales* order and its families (95% of assigned sequences) was found in soils of a permafrost thaw gradient in northern Sweden (Emerson et al., 2018). Our RNA-based approach now adds evidence that the *Caudovirales* are not only the most abundant, but also the most metabolically active members of the soil virome, contributing 97 % of the detected viral transcription activity in our dataset.

Interestingly, the activity of soil bacteria and their putative phages did not co-decline across the elevational gradient. While there is little knowledge on co-activity patterns, previous studies reported correlation between bacterial and phage abundance in marine and soil environments (Weinbauer, 2004; Williamson et al., 2017; Wommack & Colwell, 2000). For example, a meta-analysis of soil viral datasets, revealed viral abundance to be significantly positively correlated with bacterial abundance (Williamson et al., 2017). Such an abundance correlation might be explained by the dependency of phage replication on host availability, where a high bacterial host availability is expected to increase phage abundances (Williamson et al., 2017). In our study, moving towards an activity-based framework to capture dynamics of bacteria and phage communities, we found distinct patterns between the two groups. Here, complex predator-prey dynamics, such as negative feedback loops, host switching to maintain similar levels of activity across the gradient, or an increased virulence withing the declining population of the hosts might be possible explanations of the divergent pattern between bacteria and phage activity, in comparison to abundance-based studies (Breitbart et al., 2018; Trubl et al., 2018). Disentangling these mechanisms will be possible to be tested with increased sample sizes and fine-scale temporal replication, accounting for microsite conditions.

Considering both bacterial and viral co-occurrence patterns across elevational gradients could provide a template for answering questions regarding the diversity, distribution, dynamics, and interactions of viruses with their hosts and their abiotic environment (Sommers et al., 2021). The integration of an ecological framework in viral metagenomics and -transcriptomics could broadly expand our knowledge on ecosystem-level effects of viruses (Roux et al., 2021; Sommers et al., 2021).

## Conclusion

Our study provides a first glimpse on the activity of bacteria and their viruses across an elevational gradient, by assessing bacterial and viral activities with a metatranscriptomics approach. It remains unclear what the consequences are of the proportionally increased activity of viruses compared to the activity of their bacterial hosts at high elevations, and how this influences ecosystem functioning provided by bacteria, such as carbon degradation. One emerging hypothesis to be tested from our study is if bacteria produce proportionally more phages at higher elevations, because phages are more successful in replication than bacteria under harsh environmental conditions (Heilmann et al., 2010; Vos et al., 2009). This would mean that bacterial activity is increasingly turned into phage production at environmentally harsher higher elevations. To test this hypothesis, in the future metatranscriptomics could be combined with bacterial and viral abundance estimations, and with fine-scale temperature and soil property measurements. Phage exclusion experiments in the field might also contribute to gain a mechanistic understanding of the interaction between bacteria and their viruses, as well as the importance of viral activity for ecosystem processes.

## Data Accessibility

Raw sequence reads and assembled contigs will be deposited in the Sequence Read Archive under the BioProject XXXXXX

## Acknowledgments

We thank Damian Baranski for help with laboratory procedures, and Christoph Sinai and Tilman Schell (Frankfurt am Main) for support with bioinformatics.

## Authors’ contributions

D.M., and M.B. conceived the ideas; D.M. and ELN collected the data, D.M. performed laboratory work; D.M. and AS analyzed data, F.D.G. provided analytical guidance; D.M. and M.B wrote the manuscript. All authors contributed to the various drafts and gave final approval for publication.

