## Supplementary for "Elevation drives activity of soil bacteria, but not of bacterial viruses"

**Supplement**

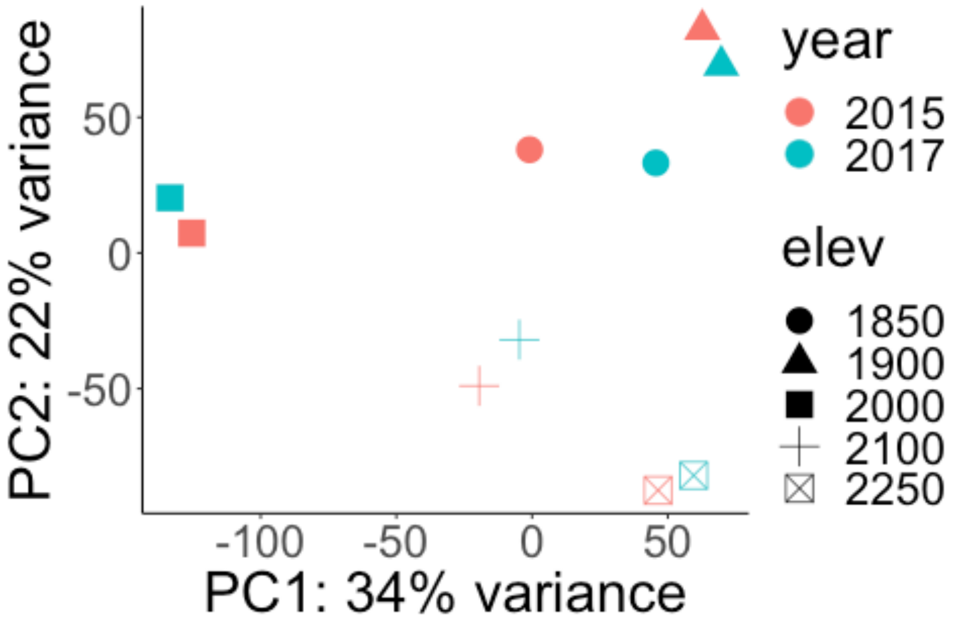


*Figure S1: Differential expression analysis revealed no significant differences in the overall expression of genes between the two sampling years.*

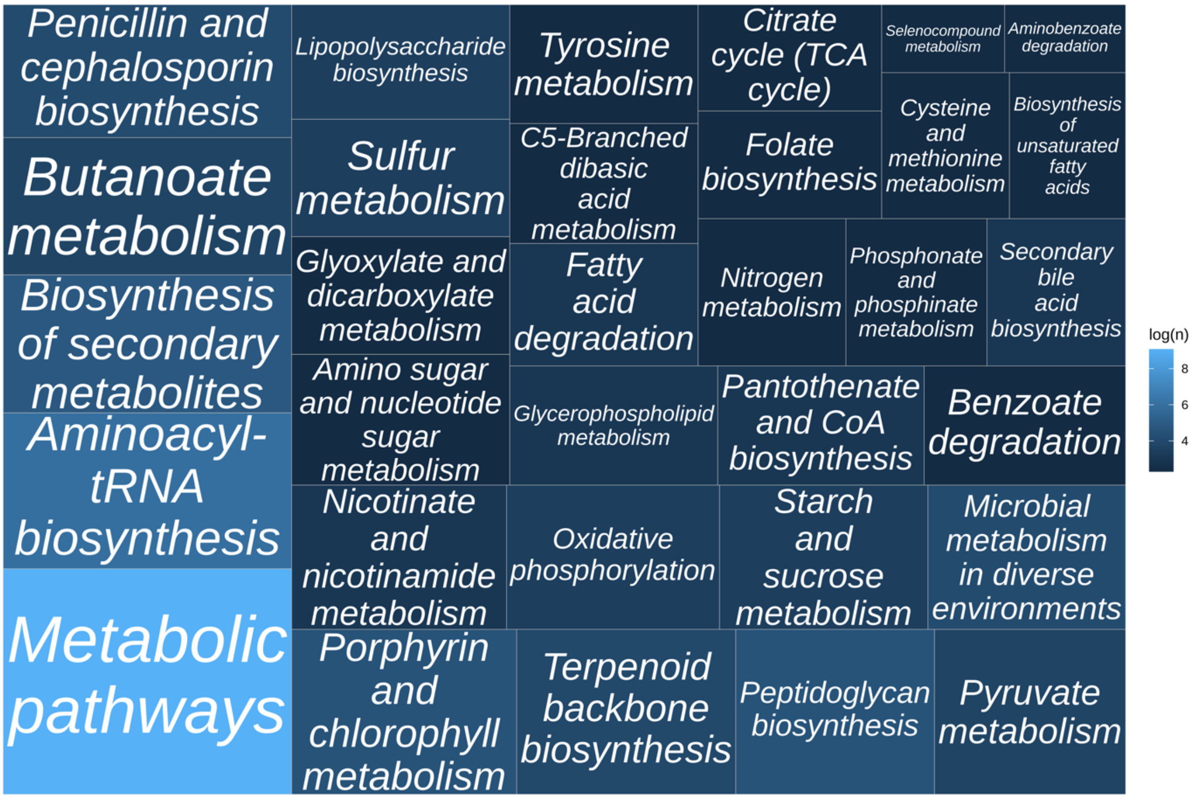


*Figure S2: Functional diversity of bacterial pathways. Color shading (dark blue to light blue) and size of square indicates proportional log-transformed activity of respective pathways.*

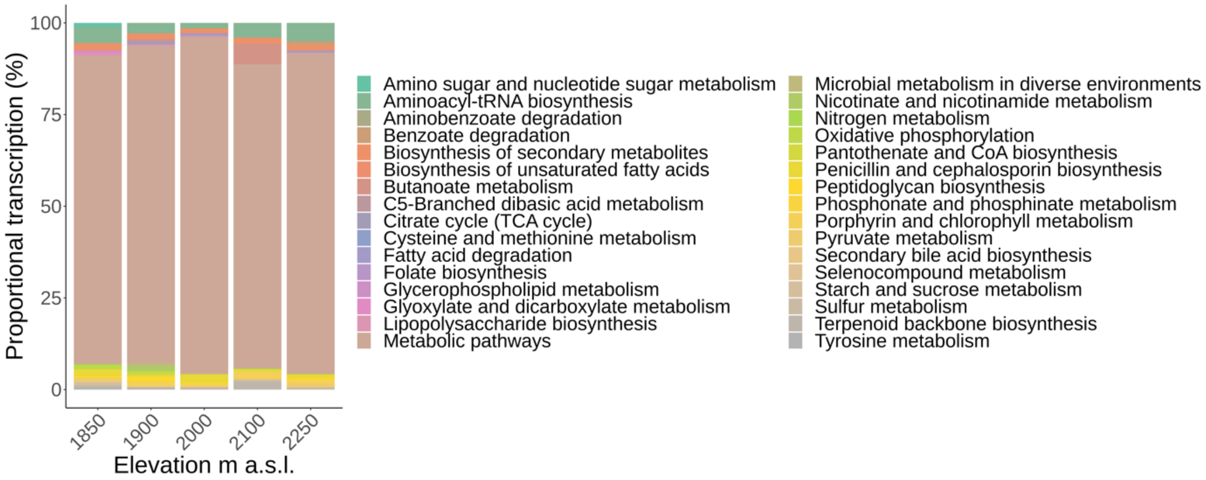


*Figure S3: Proportional transcription of bacterial pathways across the elevational gradient*

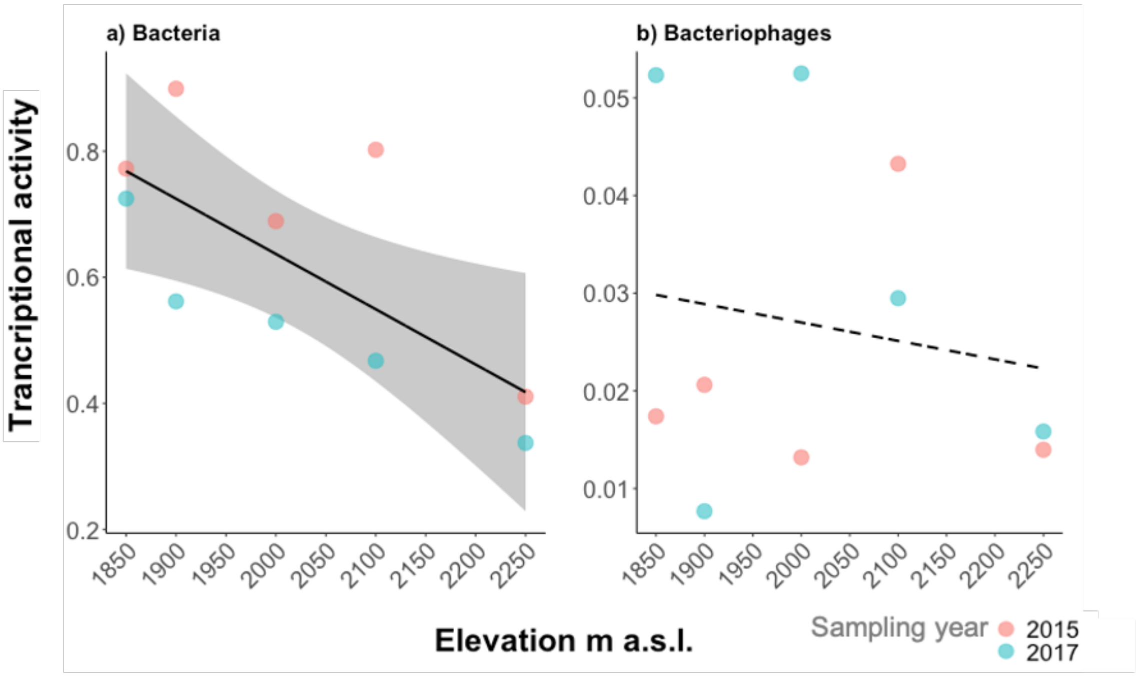


*Figure S4: Transcriptional activity normalized by mean genome size. Normalized transcriptional activity of soil bacteria significantly declined with increasing elevation. Normalized transcriptional activity of bacteriophages was not significantly affected by elevation.*

*Table S1: Sampling site characteristics.*

| Sampling ID | Elevation m a.s.l | Latitude | Longitude | Sampling year |
|---|---|---|---|---|
| S 1850 2015 | 1850 | 46.73293523 | 9.846696009 | 2015 |
| S 1850 2017 | 1850 | 46.73293523 | 9.846696009 | 2017 |
| S 1900 2015 | 1900 | 46.73260568 | 9.84903106 | 2015 |
| S 1900 2017 | 1900 | 46.73260568 | 9.84903106 | 2017 |
| S 2000 2015 | 2000 | 46.73375396 | 9.85181554 | 2015 |
| S 2000 2017 | 2000 | 46.73375396 | 9.85181554 | 2017 |
| S 2100 2015 | 2100 | 46.73468818 | 9.853815454 | 2015 |
| S 2100 2017 | 2100 | 46.73468818 | 9.853815454 | 2017 |
| S 2250 2015 | 2250 | 46.73637786 | 9.857283009 | 2015 |
| S 2250 2017 | 2250 | 46.73637786 | 9.857283009 | 2017 |

*Table S2: Normalized read counts of bacterial phyla in decreasing order.*

| Phylum | Normalized read counts |
|---|---|
| Proteobacteria | 22155055.53 |
| Actinobacteria | 4183279.34 |
| Firmicutes | 1581643.84 |
| Acidobacteria | 1070837.20 |
| Bacteroidetes | 376247.67 |
| Planctomycetes | 319830.52 |
| Chloroflexi | 173546.32 |
| Verrucomicrobia | 169267.40 |
| Gemmatimonadetes | 128310.80 |
| Armatimonadetes | 59203.89 |
| Deinococcus-Thermus | 56203.74 |
| Thermotogae | 41311.61 |
| Chlamydiae | 26589.31 |
| Euryarchaeota | 23539.60 |
| Cyanobacteria | 15642.63 |
| Spirochaetes | 8592.85 |
| Fusobacteria | 8251.88 |
| Thermodesulfobacteria | 5960.48 |
| Nitrospirae | 4056.25 |
| Elusimicrobia | 3515.72 |
| Aquificae | 2442.70 |
| Thaumarchaeota | 2188.07 |
| Lentisphaerae | 1426.40 |
| Crenarchaeota | 786.36 |
| Deferribacteres | 781.08 |
| Tenericutes | 652.69 |
| Chlorobi | 542.79 |
| Ignavibacteriae | 490.04 |
| Calditrichaeota | 389.52 |
| Abditibacteriota | 311.44 |
| Fibrobacteres | 240.83 |
| Nitrospinae | 176.93 |
| Synergistetes | 103.48 |
| Balneolaeota | 69.63 |
| Rhodothermaeota | 65.20 |
| Kiritimatiellaeota | 36.59 |
| Dictyoglomi | 35.22 |

*Table S3: Normalized read counts of phage in decreasing order.*

| **Family** | **Normalized read counts** |
|---|---|
| *Siphoviridae* | 8528.60 |
| *Myoviridae* | 7447.21 |
| *Leviviridae* | 464.69 |
| *Podoviridae* | 425.19 |
| *Autographiviridae* | 318.78 |
| *Demerecviridae* | 219.05 |
| *Drexlerviridae* | 184.53 |
| *Herelleviridae* | 147.43 |
| *Microviridae* | 118.09 |

*Table S4: Normalized read counts of bacteriophages.*

| Sampling ID | Autographi-viridae | Demerec-viridae | Drexler-viridae | Herelle-viridae | Levi-viridae | Micro-viridae | Myo-viridae | Podo-viridae | Sipho-viridae |
|---|---|---|---|---|---|---|---|---|---|
| S 1850 2015 | 0.00 | 0.00 | 51.10 | 0.00 | 0.00 | 0.00 | 343.93 | 73.28 | 649.70 |
| S 1850 2017 | 0.00 | 107.25 | 0.00 | 91.10 | 0.00 | 95.94 | 837.10 | 0.00 | 1321.60 |
| S 1900 2015 | 17.91 | 55.02 | 60.04 | 0.00 | 464.69 | 22.15 | 465.52 | 87.68 | 464.94 |
| S 1900 2017 | 0.00 | 0.00 | 0.00 | 36.22 | 0.00 | 0.00 | 64.83 | 48.17 | 319.44 |
| S 2000 2015 | 45.01 | 8.58 | 21.22 | 20.11 | 0.00 | 0.00 | 457.05 | 32.09 | 382.92 |
| S 2000 2017 | 0.00 | 0.00 | 0.00 | 0.00 | 0.00 | 0.00 | 1837.23 | 0.00 | 1999.85 |
| S 2100 2015 | 242.33 | 0.00 | 0.00 | 0.00 | 0.00 | 0.00 | 2343.14 | 0.00 | 991.08 |
| S 2100 2017 | 0.00 | 0.00 | 0.00 | 0.00 | 0.00 | 0.00 | 378.73 | 129.55 | 1274.73 |
| S 2250 2015 | 13.52 | 20.99 | 52.17 | 0.00 | 0.00 | 0.00 | 259.69 | 54.43 | 494.20 |
| S 2250 2017 | 0.00 | 27.21 | 0.00 | 0.00 | 0.00 | 0.00 | 459.97 | 0.00 | 630.15 |

*Table S5: Normalized read counts of bacteria phyla (A-B).*

| Sample ID | Abditibacteriota | Acidobacteria | Actinobacteria | Aquificae | Armatimonadetes | Bacteroidetes | Balneolaeota |
|---|---|---|---|---|---|---|---|
| S 1850 2015 | 0.00 | 50667.34 | 396173.65 | 57.52 | 9242.06 | 23472.60 | 0.00 |
| S 1850 2017 | 0.00 | 173107.44 | 646523.87 | 772.08 | 5292.09 | 89201.67 | 0.00 |
| S 1900 2015 | 200.82 | 22704.32 | 748730.46 | 354.63 | 1061.58 | 124202.93 | 39.51 |
| S 1900 2017 | 20.11 | 336166.33 | 439720.03 | 302.30 | 7219.93 | 14649.98 | 0.00 |
| S 2000 2015 | 0.00 | 41787.95 | 191758.75 | 216.86 | 1406.49 | 14591.49 | 0.00 |
| S 2000 2017 | 0.00 | 159274.68 | 377637.71 | 0.00 | 7707.34 | 17342.65 | 0.00 |
| S 2100 2015 | 0.00 | 73049.47 | 347026.07 | 0.39 | 8127.58 | 35651.53 | 0.00 |

| | | | | | | | |
|---|---|---|---|---|---|---|---|
| S 2100 2017 | 73.41 | 67776.54 | 372626.80 | 284.82 | 10358.58 | 14113.70 | 0.00 |
| S 2250 2015 | 17.10 | 88362.92 | 458991.85 | 305.66 | 1972.34 | 17984.14 | 30.12 |
| S 2250 2017 | 0.00 | 57940.20 | 204090.15 | 148.46 | 6815.90 | 25036.98 | 0.00 |

*Table S6: Normalized read counts of bacteria phyla (C).*

| Sample ID | Calditrichaeota | Chlamydiae | Chlorobi | Chloroflexi | Crenarchaeota | Cyanobacteria |
|---|---|---|---|---|---|---|
| S 1850 2015 | 0.00 | 3190.57 | 0.00 | 14965.21 | 1.47 | 1153.83 |
| S 1850 2017 | 0.00 | 10767.60 | 158.43 | 19048.75 | 538.05 | 1293.63 |
| S 1900 2015 | 23.38 | 953.15 | 23.21 | 17429.00 | 0.00 | 1224.70 |
| S 1900 2017 | 119.04 | 1051.05 | 148.87 | 28966.40 | 107.90 | 2786.62 |
| S 2000 2015 | 30.98 | 964.06 | 0.00 | 5342.04 | 31.96 | 1379.92 |
| S 2000 2017 | 0.00 | 582.11 | 0.00 | 15435.78 | 0.00 | 620.90 |
| S 2100 2015 | 114.17 | 2864.11 | 0.00 | 7360.31 | 60.60 | 1985.32 |
| S 2100 2017 | 63.13 | 2038.02 | 85.23 | 16629.46 | 0.00 | 2182.48 |
| S 2250 2015 | 38.83 | 1999.44 | 15.31 | 15116.74 | 46.39 | 1205.10 |
| S 2250 2017 | 0.00 | 2179.19 | 111.74 | 33252.63 | 0.00 | 1810.14 |

*Table S7: Normalized read counts of bacteria phyla (D-F).*

| Sample ID | Deferribacteres | Deinococcus-Thermus | Dictyoglomi | Elusimicrobia | Euryarchaeota | Fibrobacteres | Firmicutes | Fusobacteria |
|---|---|---|---|---|---|---|---|---|
| S 1850 2015 | 18.68 | 7409.53 | 0.00 | 52.74 | 1205.24 | 0.00 | 133416.88 | 387.98 |
| S 1850 2017 | 243.50 | 13441.32 | 0.00 | 161.74 | 2824.21 | 80.56 | 393817.19 | 6156.29 |
| S 1900 2015 | 28.49 | 4901.11 | 0.00 | 40.86 | 802.31 | 0.00 | 274282.13 | 317.15 |
| S 1900 2017 | 0.00 | 751.70 | 0.00 | 2462.55 | 3264.45 | 32.42 | 89156.36 | 398.71 |
| S 2000 2015 | 67.94 | 2494.54 | 14.30 | 100.19 | 1236.30 | 10.70 | 92291.26 | 82.70 |
| S 2000 2017 | 0.00 | 3893.71 | 0.00 | 240.75 | 1151.30 | 0.00 | 78846.45 | 0.00 |

| | | | | | | | | |
|---|---|---|---|---|---|---|---|---|
| S 2100 2015 | 0.00 | 10021.51 | 0.00 | 0.00 | 9059.65 | 0.00 | 119878.29 | 382.79 |
| S 2100 2017 | 0.00 | 5275.62 | 0.00 | 344.25 | 2401.70 | 0.00 | 110050.12 | 117.76 |
| S 2250 2015 | 365.63 | 3882.53 | 20.92 | 39.11 | 210.71 | 42.97 | 221086.74 | 94.35 |
| S 2250 2017 | 56.85 | 4132.18 | 0.00 | 73.52 | 1383.73 | 74.18 | 68818.42 | 314.16 |

*Table S8: Normalized read counts of bacteria phyla (G-P).*

| Sample ID | Gemma-timonadetes | Ignavibacteriae | Kiritimatiellaeota | Lentisphaerae | Nitrospinae | Nitrospirae | Planctomycetes | Proteobacteria |
|---|---|---|---|---|---|---|---|---|
| S 1850 2015 | 939.80 | 32.10 | 0.00 | 1.05 | 0.00 | 248.32 | 28431.89 | 2851419.14 |
| S 1850 2017 | 18448.46 | 0.00 | 0.00 | 0.00 | 0.00 | 321.97 | 35539.82 | 2517682.13 |
| S 1900 2015 | 1992.70 | 26.60 | 0.00 | 16.38 | 11.03 | 160.10 | 12489.59 | 3371258.35 |
| S 1900 2017 | 72420.38 | 134.39 | 12.28 | 68.86 | 82.18 | 1517.80 | 57956.60 | 1766730.17 |
| S 2000 2015 | 509.74 | 109.97 | 24.31 | 161.16 | 0.00 | 178.23 | 7488.05 | 2579795.89 |
| S 2000 2017 | 15.68 | 0.00 | 0.00 | 0.00 | 0.00 | 998.80 | 25000.87 | 1858641.93 |
| S 2100 2015 | 2645.25 | 0.00 | 0.00 | 808.47 | 0.00 | 183.69 | 25189.07 | 2953228.79 |
| S 2100 2017 | 8992.38 | 137.84 | 0.00 | 0.00 | 65.74 | 0.00 | 38605.46 | 1667136.47 |
| S 2250 2015 | 15969.18 | 49.14 | 0.00 | 8.53 | 17.99 | 184.95 | 37136.53 | 1439819.02 |
| S 2250 2017 | 6377.22 | 0.00 | 0.00 | 361.93 | 0.00 | 262.40 | 51992.65 | 1149343.65 |

*Tabelle S9: Normalized read counts of bacteria phyla (G-P).*

| Sample ID | Rhodo-thermaeota | Spirochaetes | Synergistetes | Thaumarchaeota | Thermo-desulfobacteria | Thermotogae | Tenericutes | Verrucomicrobia |
|---|---|---|---|---|---|---|---|---|
| S 1850 2015 | 0.00 | 426.51 | 0.00 | 101.42 | 432.12 | 4551.89 | 0.00 | 8325.00 |
| S 1850 2017 | 0.00 | 4861.91 | 0.00 | 162.74 | 1510.46 | 12349.46 | 220.41 | 9565.65 |
| S 1900 2015 | 29.14 | 1767.33 | 0.00 | 63.92 | 1073.52 | 5631.30 | 33.43 | 7681.69 |
| S 1900 2017 | 0.00 | 391.80 | 0.00 | 699.64 | 433.83 | 4505.78 | 22.33 | 46894.64 |

| | | | | | | | | |
|---|---|---|---|---|---|---|---|---|
| S 2000 2015 | 36.06 | 251.37 | 30.61 | 150.44 | 188.00 | 1909.40 | 4.33 | 7194.51 |
| S 2000 2017 | 0.00 | 0.00 | 0.00 | 700.23 | 0.00 | 386.31 | 0.00 | 24769.32 |
| S 2100 2015 | 0.00 | 339.81 | 0.00 | 0.00 | 0.00 | 3692.59 | 0.00 | 13776.22 |
| S 2100 2017 | 0.00 | 27.04 | 0.00 | 153.64 | 1850.56 | 2802.90 | 157.25 | 9476.02 |
| S 2250 2015 | 0.00 | 260.83 | 0.00 | 29.81 | 351.94 | 3521.33 | 187.67 | 11928.33 |
| S 2250 2017 | 0.00 | 266.25 | 72.87 | 126.22 | 120.05 | 1960.65 | 27.25 | 29656.01 |

*Table S10: Mean genome sizes of bacterial phyla in base pairs (bp).*

| **Phylum** | **Mean genome size (bp)** |
|---|---:|
| Acidobacteria | 5658470 |
| Aquificae | 1661868 |
| Armatimonadetes | 5788381 |
| Chlamydiae | 1509233 |
| Chloroflexi | 5186494 |
| Deferribacteres | 2821942 |
| Deinococcus-Thermus | 3547675 |
| Fibrobacteres | 3064425 |
| Fusobacteria | 2240880 |
| Gemmatimonadetes | 6360666 |
| Kiritimatiellaeota | 6334608 |
| Lentisphaerae | 5372493 |
| Planctomycetes | 7004266 |
| Proteobacteria | 3820901 |
| Synergistetes | 2877858 |
| Tenericutes | 5658470 |
| Thermodesulfobacteria | 1990616 |
| Thermotogae | 2183525 |
| Verrucomicrobia | 4725469 |

*Table S11: Mean genome sizes of viral families in base pairs (bp).*

| **Family** | **Mean genome size (bp)** |
|---|---:|
| Autographiviridae | 41586 |
| Demerecviridae | 113454 |
| Drexlerviridae | 49426 |
| Herelleviridae | 147732 |
| Leviviridae | 3747 |
| Microviridae | 5033 |
| Myoviridae | 125703 |
| Podoviridae | 55839 |
| Siphoviridae | 52792 |
